# Tensor decomposition for data-driven analysis of OP-MEG: a feasibility study

**DOI:** 10.64898/2026.11.20.766579

**Authors:** Hugo Wharton, Arjun Ramaswamy, Stephanie Mellor, Sahitya Puvvada, Matthew Walker, Daniel Bush, Tim Tierney, Umesh Vivekananda, Mehrnoosh Sadrzadeh

**Affiliations:** Department of Physics & Astronomy, University College London, London, United Kingdom; Quantum Learning Labs, Department of Computer Science, University College London, London, United Kingdom; Department of Imaging Neuroscience, UCL Queen Square Institute of Neurology, University College London, London, United Kingdom; Research Department of Epilepsy, UCL Queen Square Institute of Neurology, University College London, London, United Kingdom; National Hospital for Neurology and Neurosurgery, University College London Hospitals NHS Foundation Trust, London, United Kingdom; Institute of Cognitive Neuroscience, University College London, London, United Kingdom; Translational Neuromodeling Unit, Institute for Biomedical Engineering, University of Zurich & ETH Zurich, Zurich, Switzerland

**Keywords:** OP-MEG, tensor decomposition, canonical polyadic decomposition, source reconstruction

## Abstract

Optically Pumped Magnetometer magnetoencephalog-raphy (OP-MEG) enables flexible, close-to-scalp recordings unlocking access to new patient cohorts and naturalistic paradigms. The data collected during OP-MEG sessions is naturally multi-dimensional, often containing information across patients, trials, conditions, space, frequency and time which can be stored in tensors. Canonical polyadic decomposition (CPD) is an unsupervised method that separates multidimensional data into components with linked spatial, spectral and temporal profiles. This allows neural responses to be characterised across brain regions, frequency and time simultaneously, without specifying these features in advance. This makes CPD well suited to OP-MEG data, where signals naturally vary across these dimensions. Here, we applied CPD to whole-brain, source-reconstructed OP-MEG data from three participants with epilepsy during a simple visuomotor task, modelling neural activity jointly across brain regions, frequency and time. We split these subject-specific tensors into event-related power decrease and increase (referred to as ERD and ERS for convenience) tensors so that a more interpretable and computationally efficient non-negative CPD variant could be applied. CPD paired with consensus clustering on spatio-spectral-temporal networks successfully identified cross-patient, simultaneous motor beta and visual alpha suppression. Permutation-based null distribution tests were used to exclude tensors that lacked significant low-rank structure. These findings demonstrate the applicability of tensor decompositions to electrophysio-logical data as the pipeline recovers canonical visuomotor responses without prespecifying region, band or latency.

## Introduction

Optically pumped magnetometer based MEG (OP-MEG) is opening up a wide range of paradigms, including naturalistic tasks (Gibson et al., 2026; Spedden et al., 2025; O’Neill et al., 2025), mobile systems (M. Brookes et al., 2026) and clinical applications such as mesial temporal lobe epilepsy studies and surgery planning (Feys et al., 2025; Mellor et al., 2024). These settings bring their own analysis challenges: sensor layouts may differ across participants and naturalistic paradigms frequently rule out time-locking across participants. Source reconstruction addresses the first of these by mapping heterogeneous sensor-level data into a common anatomical space, but the subsequent analysis must still accommodate the latter.

Conventional M/EEG analyses require the region, frequency band and latency of interest to be specified in advance, which is restrictive in exploratory, naturalistic and clinical studies where these are rarely known a priori. Data-driven methods remove this requirement, but to be practical for researchers and clinicians, who typically work under time and funding constraints, they must be interpretable, statistically grounded and computationally inexpensive. Deep learning (LeCun et al., 2015) rarely meets these criteria in applied neuroimaging: it is costly to train, has a high barrier to entry and, more fundamentally, produces models that are difficult to interpret (Rudin, 2019; Haufe et al., 2014). Many studies therefore rely on classical methods such as principal component analysis (PCA) or hidden Markov models (HMMs) (Vidaurre et al., 2017; Gohil et al., 2024). These operate on matrices, so multidimensional data such as signals that vary across region, frequency and time must first be reshaped before decomposition. Although the resulting components can be reshaped back afterwards, the decomposition itself is fitted after dimensions have been combined and therefore does not explicitly model spatial, spectral and temporal profiles as separate but linked dimensions. Tensor decomposition instead estimates these profiles jointly, allowing each component to describe a pattern in terms of where, at what frequency and when it occurs.

Canonical polyadic decomposition (CPD) is an unsupervised tensor decomposition that, like PCA, expresses data as a sum of components. Unlike PCA, CPD can, under established conditions, recover an essentially unique set of components apart from their order and overall scaling (Stegeman and Sidiropoulos, 2007). In practical terms, this reduces ambiguity in how the same underlying pattern can be represented, helping each component retain a consistent spatial, spectral and temporal interpretation. These theoretical properties motivate the choice of CPD over matrix methods, although a direct empirical comparison on OP-MEG tensors is not performed here and remains an open methodological question. CPD has been applied broadly in signal processing, machine learning and biomedical image analysis (Sedighin, 2024; Burch et al., 2025; Ballard and Kolda, 2025). Provided low-rank structure is present in the data, the resulting components are directly interpretable (Kolda and Bader, 2009): in this work, each rank-1 component carries an explicit spatial, spectral and temporal profile, which can be compared across participants. CPD can consequently supplement or replace conventional analysis pipelines, yielding whole-brain, frequency- and time-resolved descriptions of neural activity without prior specification of region, band or latency.

CPD has been applied to neuroimaging data across third-, fourth- and fifth-order tensors: to characterise EEG dynamics (Li et al., 2019), to identify neuronal activation patterns during learning in mice (Williams et al., 2018) and to perform multi-subject MEG analyses (Belyaeva et al., 2023). As a higher-order feature extraction method, CPD also yields components usable as input features, for instance to classify conditions such as epilepsy in large intracranial EEG cohorts (Wagh et al., 2025). Despite these applications, CPD has seen relatively limited use in neuroimaging, particularly in source-space MEG. This may partly reflect the additional modelling choices introduced by tensor methods, including rank selection and the alignment of components across participants. To our knowledge, CPD has not previously been applied to source-reconstructed MEG data.

In this work, baseline-relative power can reflect either decreases or increases in activity relative to baseline. We therefore analyse event-related desynchronisation (ERD) and synchronisation (ERS) separately and fit each using non-negative CPD (NN-CPD). This prevents positive and negative responses from cancelling within a component, allowing the loadings to be interpreted directly as the contribution of a region, frequency and time interval to either power suppression or enhancement. Fewer components are also required to describe each tensor, reducing the computational cost across the repeated fits required to assess solution stability.

Sensorimotor beta rhythms (14–30 Hz) are modulated across a variety of active and passive movements, characteristically exhibiting a power decrease prior to and during movement, followed by a post-movement rebound (Pfurtscheller et al., 1998; Pfurtscheller and Lopes Da Silva, 1999; Jurkiewicz et al., 2006). Similarly, occipital alpha rhythms (7–14 Hz) are strongly modulated during visual processing, typically showing ERD in response to primary visual stimuli (Pfurtscheller and Lopes Da Silva, 1999; M. J. Brookes et al., 2005). OP-MEG has been shown to detect both motor beta ERD (Stærmose et al., 2026; Boto et al., 2017) and occipital alpha and gamma rhythms (Zhang et al., 2020; Rhodes et al., 2024; Iivanainen et al., 2019). These responses therefore provide physiological benchmarks for the proposed analysis: their recovery shows that an unsupervised whole-brain decomposition can isolate established task-related responses in OP-MEG without defining the relevant region, frequency band or latency in advance.

Here, OP-MEG recordings were acquired from three participants with epilepsy during the visuomotor section of a reward learning paradigm (Ramaswamy et al., 2026), and NN-CPD was applied to source-reconstructed data epoched around choice events. Each participant’s tensor was decomposed independently, rather than stacked into a single group tensor, and shared patterns were then aligned to form group-level clusters. This design allows a mode such as time to be relaxed in the cross-participant analysis, which is necessary for paradigms where time-locking across participants is not possible. These clusters were compared with the OPM and wider neuroimaging literature.

The primary aim of this study was therefore methodological: a proof of concept presenting a tensor-based alternative to matrix methods for source-reconstructed OP-MEG. As each participant’s tensor is decomposed independently, each recording provides a separate test of whether reproducible low-rank structure is present; the three participants here are therefore treated as replications of the pipeline rather than as a sample from which population-level inferences are drawn. The analysis was evaluated at three levels. First, decomposition stability and reconstruction performance were used to assess reproducible low-rank structure. Second, temporally shifted surrogate trials tested whether this structure depended on alignment to the choice event. Third, the recovered clusters were compared with established motor beta and visual alpha responses as a test of physiological plausibility.

## Method

**Figure 1:**
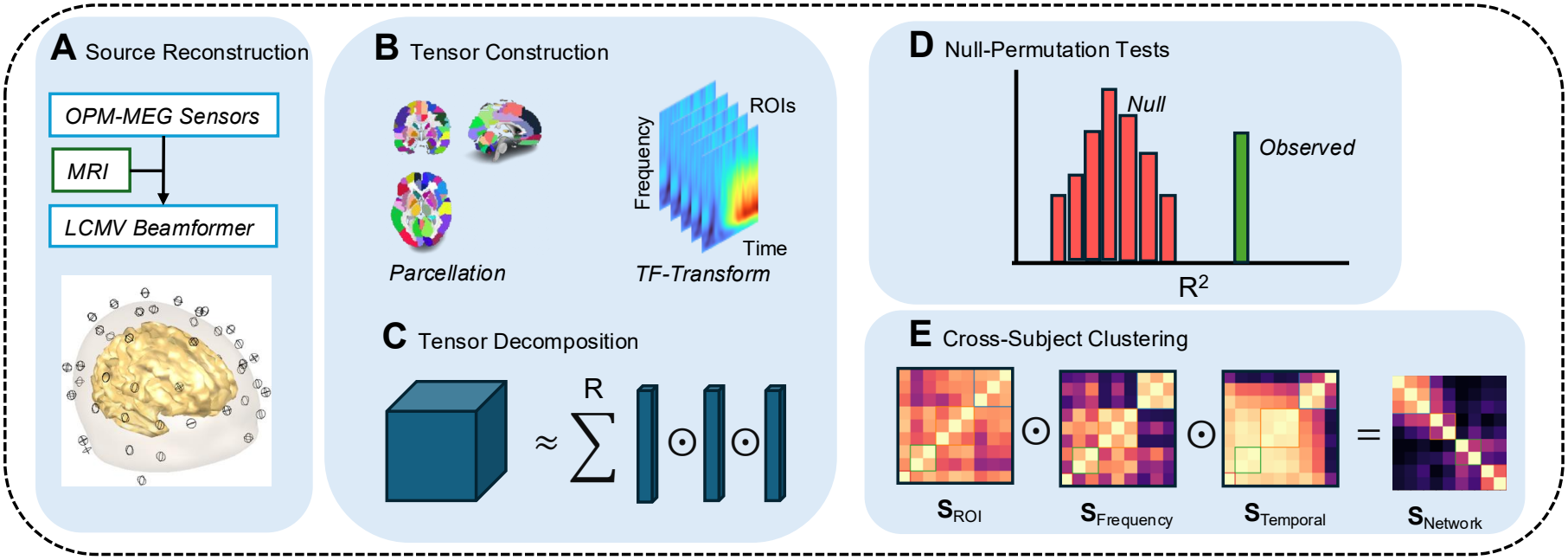
Overview of the analysis pipeline. Participant-specific OP-MEG recordings were preprocessed in sensor space, beamformer reconstructed onto a common source-space parcellation and represented as region-by-frequency-by-time tensors. ERD and ERS tensors were decomposed independently for each participant using NN-CPD, after which components were aligned across participants using constrained consensus clustering. R = Number of Components.

### Data Collection and Experimental Setup

Bespoke 3D-printed helmets, designed from each participant’s MRI, housed 23/23/25 triaxial and 28/34/32 dual-axis QuSpin sensors (participants 1–3 respectively), providing full head coverage (Figure 2). Data were recorded from three participants with epilepsy during a reward learning paradigm (Ramaswamy et al., 2026), time-locked to the button press events in which participants selected an option, after which their choice was displayed onscreen. Ethical approval was given by the Essex Research Ethics Committee in the UK (REC Reference 18/EE/0220) and informed consent obtained. Participant demographics, valid trial counts and recording lengths are provided in Appendix Table A1.

**Figure 2:**
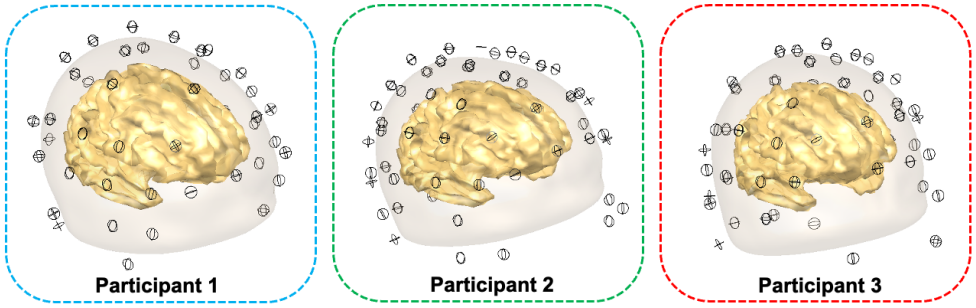
Sensor layouts for participants 1–3, showing the distribution of OP-MEG sensors across the head. Each participant wore an individually designed helmet providing whole-head coverage.

Each trial therefore combines a motor/sensory event (the button press) with subsequent visual processing of the presented stimulus.

### Sensor Space Preprocessing

Sensor-space preprocessing was performed in SPM (Tierney et al., 2025). Data were downsampled to 1000 Hz, and bad channels identified from power-spectral-density plots were manually removed. 4th and 5th order Butterworth filters were applied: a 1 Hz high-pass, a 48–52 Hz notch, and a 70 Hz low-pass. Adaptive Multipole Modelling (AMM) (Tierney et al., 2024) was used to shield against environmental noise; the resulting covariance rank-deficiency later mitigated via 5% Tikhonov regularisation.

Continuous OP-MEG sensor data was epoched into 2.5 s bins ([−1000, +1500] ms about the per-trial choice-selection button press) for stable decomposition estimates, rejecting trials with ambiguous button-press timings (e.g due to multiple presses) or excessive channel variance. Each participant’s OP-MEG sensor geometry was co-registered to their anatomical MRI for source reconstruction.

### Source Reconstruction

Source reconstruction links two spatial domains: the sensor space of the raw OP-MEG recordings and the source space in which neural activity is interpreted by the tensor decomposition. An LCMV beamformer (Van Veen and Buckley, 1988; Westner et al., 2022) projected the sensor-level data into source space, yielding an estimate of the dipole moment at location *r* and time point *t* throughout the brain. Spatial filters were computed on a 5 mm source space grid paired with a single-shell head model derived from each participant’s individual MRI. Source orientation at each grid point was fixed to the direction of maximum signal strength via SVD, and the resulting source estimates were warped to MNI, a common geometrical space. As all participants had whole-head coverage, a whole-brain analysis was appropriate.

Subsequent analysis was restricted to the 70 cortical ROIs (Table A2) since the purpose was to assess the feasibility of the pipeline on well established cortical visual/motor responses in areas of high MEG sensitivity.

### Tensor Construction

Spectral information was extracted from each trial’s ROI time series by wavelet convolution (Lina et al., 2014), yielding 1 Hz bins from 4–70 Hz at 0.01 s steps. Trial spectra were trimmed to [−200, +1000] ms, lognorm baselined, and averaged to obtain the induced response (evoked activity subtracted), giving a third-order *X ∈* ℝ^ROI*×*Freq*×*Time^ tensor per participant. A fourth-order *X ∈* ℝ^Subject*×*ROI*×*Freq*×*Time^ tensor was trialled but discarded since a shared subject CPD forces every component to exist in every participant with only a scalar difference, given the heterogeneity of the epileptic cohort and the variability in SNR due to trial count, third-order decompositions were retained. A per participant decomposition with clustering on components (Argüelles et al., 2014) permits absence of shared patterns and SNR quality variation. Because event-related synchronisation (ERS) and desynchronisation (ERD) can be treated as separable, though not independent, neurobiological processes, each tensor was split into ERS and ERD components. This split permits a non-negative tensor decomposition, which yielded greater stability and model fit with fewer components and lower computational cost.

The participant-specific formulation also relaxes the assumption imposed by a joint subject-by-region-by-frequency-by-time model that each component has a corresponding expression in every participant. Components can instead remain participant-specific and are only assigned to a group-level cluster when their spatial, spectral and temporal profiles are sufficiently similar.

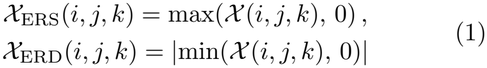

### Non-negative CP Tensor Decomposition

Although a third-order tensor can be unfolded into a matrix, this discards the higher-order multilinear structure, making tensor methods well suited to neuroimaging data with naturally distinct spatial, spectral and temporal modes. Canonical Polyadic Decomposition (CPD) approximates a tensor as a sum of *R* rank-one components, each an outer product of *N* vectors, computed via the following formulae, where *λ*_*r*_ weights component *r*:

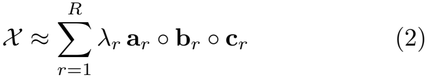

The above formula can equivalently be stated in factor-matrix form as *X ≈* **[A, B, C**]], where **A, B**, and **C**, are in 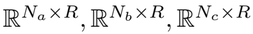 respectively and [*·*] represents the Kruskal operator, summing the weighted outer products of the factor-matrix columns. A non-negativity constraint was imposed on all modes, and the NN-CPD was computed by Alternating Least Squares (ALS) (Kolda and Bader, 2009).

We imposed non-negativity on the factor matrices because signed CPD fitted the tensors poorly and needed a computationally expensive rank (*R ≥* 10). The subsequent statistical tests rely on repeated model runs, so we prioritised computational efficiency. Discussions of model rank selection can be found in the appendix.

**Figure 3:**
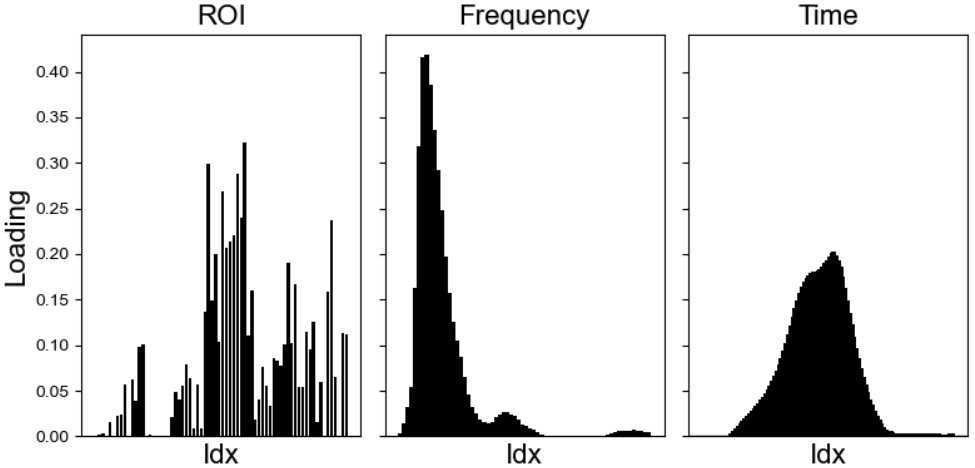
Example NN-CPD component from participant 1. The spatial, spectral and temporal loading vectors jointly define one rank-one pattern in the baseline-normalised ERD tensor.

### Monte-Carlo Permutation Simulations

Permutation null test statistics were collected to assess whether the distribution of trials showed significant effects and that the CPD model was not fitting to noisy, simulated data at the chosen rank. The per participant observed ERD/ERS tensors were decomposed to obtain the baseline test statistic, chosen as *R*^2^. To construct an empirical null distribution, random circular shifts were applied [*N*_Sim_ = 500] times along the temporal axis of individual trials, disrupting task-locked temporal alignment whilst preserving the inherent spectral and spatial covariance. These permuted trials were subsequently baseline-normalised and averaged to generate surrogate third-order tensors. Decomposition of these null tensors at the matched rank yielded a distribution from which uncorrected exact right-tailed p-values were derived (Appendix: Figure A1).

### Cross-participant Clustering

Components from each participant’s ERD and ERS decompositions (R = 3) were grouped by network similarity,

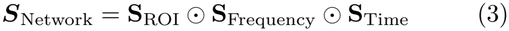

Here **S**_**ROI**_ and **S**_**Freq**_ are cosine similarities between loading vectors, and **S**_**Time**_ is the maximum normalised cross-correlation within *±*100 ms, allowing for interparticipant latency differences. The product requires agreement in all three modes. Clusters were obtained by constrained clustering (Grötschel and Wakabayashi, 1989), solved as a binary integer program maximising within-cluster similarity (HiGHS solver (Huangfu and Hall, 2018)). The program was subject to a thresh-old (*S*_*Network*_ *≥* 0.6), a cannot-link constraint (at most one component per participant per cluster) and transitivity. A sweep of thresholds was performed which showed the (0.4-0.6) range produced the same clusters shown here. At higher or lower values, components were dropped or included despite large discrepancies. For stability, the similarity matrix was perturbed with symmetric Gaussian noise (*σ* = 0.05) over 500 iterations, and the clustering was re-solved each time (Monti et al., 2003). Final clusters were obtained by clustering the resulting co-association matrix, and mean within-cluster co-association is reported as a stability score. The noise level and lag window were fixed a priori.

All preprocessing, source reconstruction, tensor decompositions, and Monte-Carlo permutation simulations (*N*_Sim_ = 500) were executed locally on a standard consumer laptop (Apple MacBook M4 Pro) without GPU acceleration.

## Results

### Time-Frequency Plots

Time frequency plots averaged over all 70 cortical ROIs acted as whole-cortex descriptive summaries; they show beta band ERD and subsequent ERS across participants with similar temporal distributions. Alpha ERD is present in participants 2 and 3 but is obscure in participant 1; the tensor decomposition and post-hoc matching solves this by introducing simultaneous spatial specificity and group level patterns whilst maintaining spectral and temporal precision.

**Figure 4:**
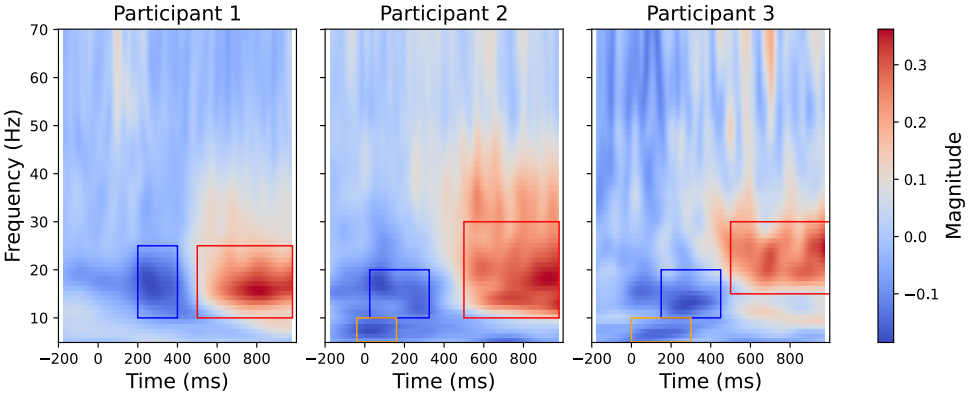
Whole-cortex time-frequency representations around button presses for each participant, computed from LCMV beamformer time series and averaged across the 70 cortical ROIs. Spatial averaging blurs some alpha and beta responses, which motivates a decomposition that keeps spatial specificity alongside time and frequency.

### Clusters

Table 1 shows the results of permutation null tests, confirming significant task-related signal 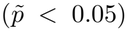 within all participants’ ERD and ERS tensors except P3 ERD which was excluded from further analyses and reflects the reduced SNR due to low trial count. The result of the simulation also showed that the CPD model was not able to fit to the null data since there was insufficient low rank structure as present in the observed data. This is a statement about task-locking, not reproducibility; decomposition stability is reported separately in the appendix. Because CPD has permutational ambiguity and hence carries no correspondence across participants, individual decompositions cannot be pooled or interpreted jointly without an explicit alignment step. Components were therefore matched across participants by constrained consensus clustering. Cross-participant clusters are presented first, with the beta-band and alpha-band clusters possibly corresponding to the motor and visual demands of the choice event, followed by the later ERS cluster. Components that did not enter any cluster are not reported: since the aim of this work is a group-level pipeline. ERD clusters are plotted in blue and ERS in red.

**Table 1:**
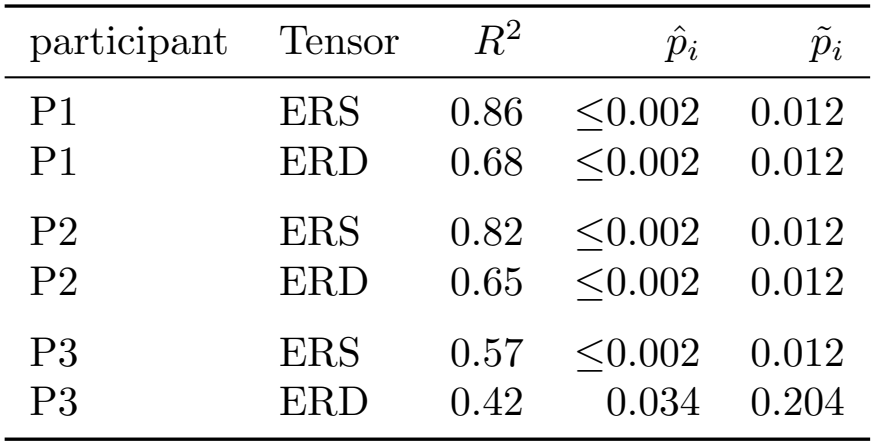
Permutation test results for six CP decompositions (three participants, two tensors each). 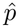 are righttailed permutation *p*-values; 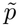 are Bonferroni-corrected across all six tests.

| participant | Tensor | R<sup>2</sup> | p̂<sub>i</sub> | p̃<sub>i</sub> |
|---|---|---|---|---|
| P1 | ERS | 0.86 | ≤0.002 | 0.012 |
| P1 | ERD | 0.68 | ≤0.002 | 0.012 |
| P2 | ERS | 0.82 | ≤0.002 | 0.012 |
| P2 | ERD | 0.65 | ≤0.002 | 0.012 |
| P3 | ERS | 0.57 | ≤0.002 | 0.012 |
| P3 | ERD | 0.42 | 0.034 | 0.204 |

### Occipital Alpha Desynchronisation

Each cluster average can be plotted as in Figure 5, where the y-axis of the spectral-temporal plots is normalised component loadings and the glass brain spatial plot shows the top 20% by loading of ROIs.

**Figure 5:**
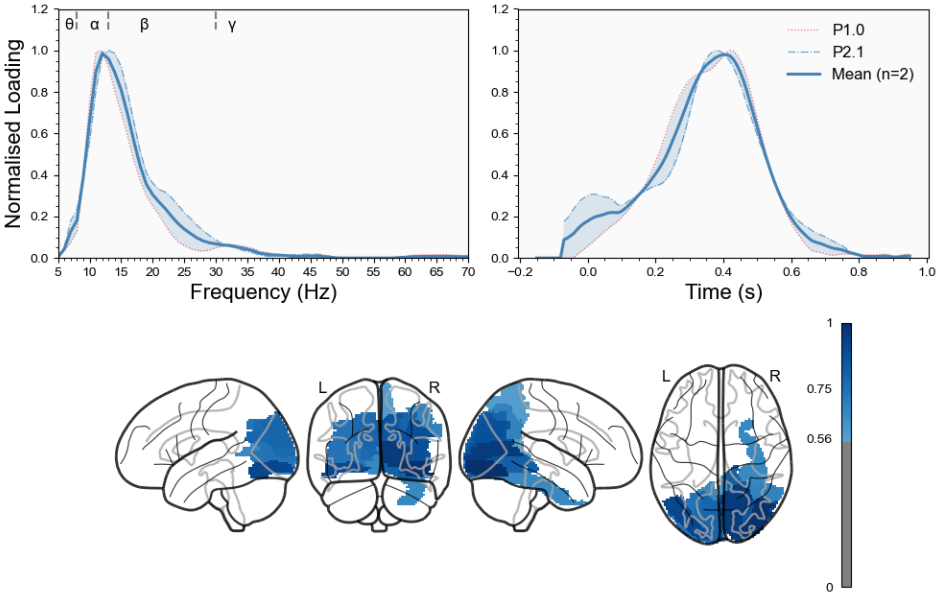
Occipital alpha ERD cluster recovered across participants 1 and 2. The component combines an alpha-band spectral profile, an occipital spatial distribution and a post-choice temporal decrease in power. Normalised loading profiles averaged are shown with error bars showing the range, regions of strong cross-subject agreement occur at the peaks.

The occipital alpha ERD cluster (Figure 5) loaded on visual parcels in both participants with a ~300-400 ms latency. No non-visual region entered the ten highest loadings in either. Participant 2 was approximately symmetric and weighted towards medial and dorsal occipital cortex (cuneus 0.283 and 0.269; inferior occipital 0.261 and 0.237; calcarine and superior occipital 0.222–0.236), whereas participant 1 was right-lateralised and weighted towards ventral and lateral occipital cortex (inferior occipital 0.323; calcarine 0.298; middle occipital 0.289; lingual 0.269). In both cases the loading was distributed across more than ten parcels rather than concentrated in two, unlike the sensorimotor beta clusters (Figure 6). This follows the expectation that lower-frequency rhythms recruit larger cortical areas and that alpha ERD is spatially less discrete than beta ERD (Pfurtscheller and Lopes Da Silva, 1999), and the confinement to visual cortex was obtained without restricting analyses spatially.

**Table 2:**
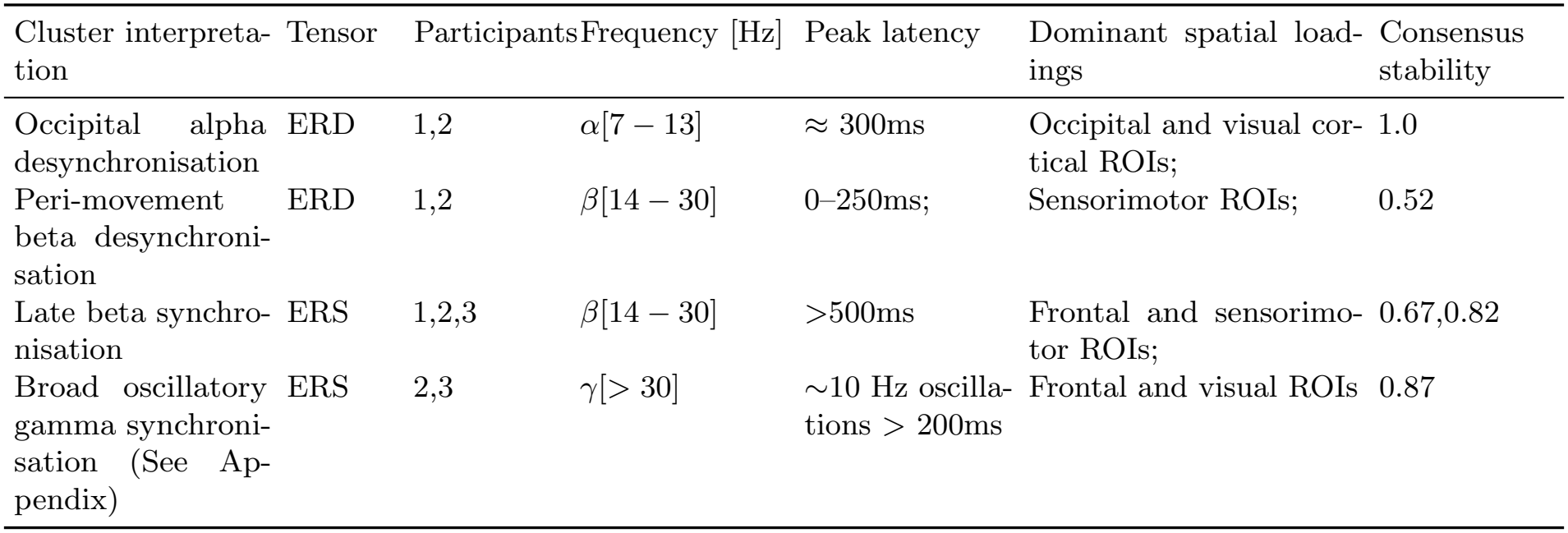
Group-level cluster inventory. Individual ROI loadings are found in the text.

| Cluster interpreta-tion | Tensor | Participants | Frequency [Hz] | Peak latency | Dominant spatial load-ings | Consensus stability |
|---|---|---|---|---|---|---|
| Occipital alpha desynchronisation | ERD | 1,2 | α[7 − 13] | ≈ 300ms | Occipital and visual cor-tical ROIs; | 1.0 |
| Peri-movement beta desynchroni-sation | ERD | 1,2 | β[14 − 30] | 0–250ms; | Sensorimotor ROIs; | 0.52 |
| Late beta synchro-nisation | ERS | 1,2,3 | β[14 − 30] | >500ms | Frontal and sensorimo-tor ROIs; | 0.67,0.82 |
| Broad oscillatory gamma synchroni-sation (See Ap-pendix) | ERS | 2,3 | γ[> 30] | ∼10 Hz oscilla-tions > 200ms | Frontal and visual ROIs | 0.87 |

**Figure 6:**
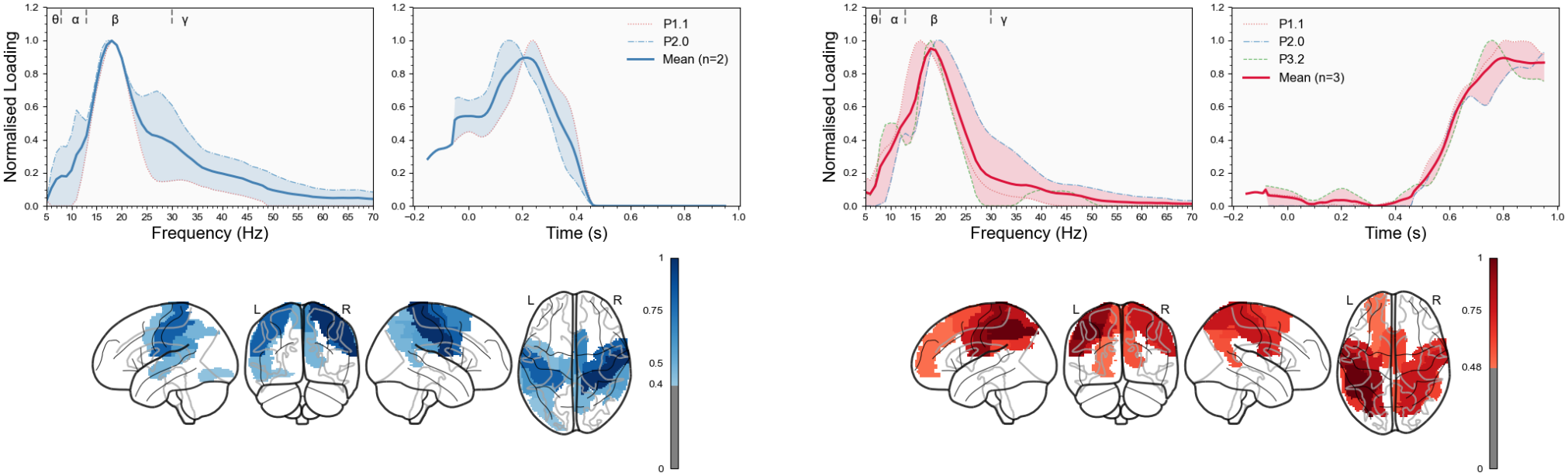
Cross-participant motor beta clusters: **A**, beta-band ERD around the button press with strongest loadings in sensorimotor regions. **B**, later beta-band ERS with frontal and sensorimotor loadings, consistent with a post-movement beta rebound. Spatial maps show the top 20% of ROI loadings; spectral and temporal profiles are normalised within component.

### Motor Cortex Beta Desynchronisation and Synchronisation

Beta ERD and subsequent ERS in the motor cortex has been reported across modalities and experimental paradigms in OP-MEG (Boto et al., 2017) and conventional MEG (Jurkiewicz et al., 2006) (Pfurtscheller et al., 1998; Pfurtscheller and Lopes Da Silva, 1999). An ERD cluster expressed in participants 1 and 2 (Figure 6) was concentrated in the beta range, peaked around the button press and showed its strongest spatial loadings in sensorimotor regions. A later ERS cluster showed beta-band synchronisation after approximately 500 ms, with frontal and sensorimotor loadings.

Components assigned to the ERD cluster loaded most strongly on postcentral gyrus (0.29 and 0.33), consistent with reports localising movement-related beta ERD to the hand region of postcentral gyrus (Jurkiewicz et al., 2006). Both peaked at ~18 Hz, below the 20–24 Hz handarea beta ERD of the example subject in (Pfurtscheller and Lopes Da Silva, 1999), whose review recommends reactive bands be determined per subject rather than fixed a priori. Here that estimate is the frequency loading vector itself, obtained without a separate band-selection step. Temporal profiles were broad and already suppressed at the −600 to −300 ms baseline, which falls within the canonical pre-movement desynchronisation window; ERD magnitude and onset are therefore compressed relative to rest (the Challenges and Limitations section).

The ERS cluster loaded on left precentral and left postcentral gyrus in all three participants, contralateral to the button press, with precentral ranked first in participants 1 and 2. All three components peaked at 15–20 Hz and rose from ~400 ms, consistent with post-movement beta synchronisation (Pfurtscheller and Lopes Da Silva, 1999). The anterior shift against the postcentral-dominant ERD cluster follows the direction reported by (Jurkiewicz et al., 2006), whose ERD and rebound peaks are separated by 16 mm and therefore below AAL2 resolution; the correspondence is descriptive rather than a source-level dissociation.

## Discussion

The tensor decomposition framework presented here serves as a proof of concept, showing that tensor methods are relevant and feasible for the analysis of source-reconstructed OP-MEG. The low-rank NN-CPD model captured reproducible, statistically significant task-locked structure in the observed tensors. The validation targets here were strongly predicted by the task design and well established in the literature; the pipeline is therefore evaluated for its capacity to reproduce known structure - a prediction for discovery of unknown structure. It also recovered well-established visuomotor alpha and beta components in a fully unsupervised manner, giving researchers an easily interpretable, data-driven alternative to hypothesis-led analyses.

Since the OP-MEG sensor layouts varied across participants, source reconstruction was used to bring the data into the AAL2 atlas, a common space in which patterns were discovered. The whole brain ROI subset selected allowed for simultaneous discovery of different spatial patterns even if their temporal and spectral profiles were highly similar since the network distance matrix requires strong agreement across all modes.

Participant tensors were decomposed individually rather than stacked into a single higher-order tensor. This keeps the components interpretable while allowing more flexibility: one mode, such as time, can be relaxed in the cross-participant analysis. That flexibility matters for many OP-MEG datasets, particularly naturalistic paradigms, where responses cannot be time-locked across participants.

### Challenges and Limitations

The main limitation is sample size; with three participants, group-level conclusions are weak and serve demonstrative purposes without more participants or healthy controls. Trial and sensor count also differed across participants. The resulting SNR differences probably affect whether a component is recovered at all and then further whether it is assigned to a cluster, although sub-cortical ROIs were excluded. The OP-MEG sensors used here lack high gamma sensitivity which limited the scope of discovery, applying the method to more recent datasets would be useful.

Decomposing trial-averaged tensors was appropriate since the aim of this study was feasibility on well-characterised responses. However, averaging discards trial-to-trial variability which a decomposition of fourth-order tensors with trial as mode would retain, although under the proposed model selection heuristics and statistical tests, the computational barrier would be significant without access to large compute facilities. There is also no ground truth on simulated data with known components.

Several of the choices made during analysis likely shape the results: the beamformer and regularisation, parcellation atlas and the decomposition rank which was fixed at R=3 for all tensors. The normalisation of factors and CPD scaling variance limits interpretation since absolute differences in power were lost. These could be preserved by joint decompositions with a shared participant mode, or by calculating difference-tensors relative to a reference participant or condition. Splitting ERD and ERS tensors permits the use of non-negative decompositions, but this sacrifices the signed structure such that coupled patterns such as the beta ERD and later ERS are recovered separately.

### Further Work

Validation on a larger cohort with complete reporting of recovered and unmatched components is the natural next step. Many adjustments could also be made to the pipeline depending on the aim of the study. For example, as discussed before, relaxing the time mode similarity during clustering could be used to analyse naturalistic paradigms where task locking is not possible. Similar spatial and spectral features could be discovered whilst the latency can be allowed to vary.

In a study where trials were maintained, bootstrapping could be used to obtain distributions on each component from which confidence intervals could be calculated and used to filter noisy components. This could enable statistically robust unsupervised pipelines in clinical or neuroscience settings.

## Conclusion

This work presented a non-negative tensor decomposition pipeline for source-reconstructed OP-MEG, in which region *×* frequency *×* time tensors were split into ERD and ERS components, decomposed independently per participant with NN-CPD, and aligned across participants by constrained consensus clustering. Five of the six decompositions contained task-locked low-rank structure exceeding a circular-shift surrogate null. Across two to three participants per effect, clusters recovered occipital alpha ERD, motor beta ERD, and post-movement beta ERS without prespecifying regions or bands. Each component carried an explicit spatial, spectral and temporal profile, so the recovered networks could be compared directly with established findings from both OP-MEG and cryogenic MEG.

With three participants, these results are a demon-stration of feasibility rather than a basis for group-level inference. Their value lies in showing that the pipeline isolates known physiological responses from whole-brain data in a fully unsupervised manner, which is the precondition for applying it where such responses are not known in advance. Because participants are decomposed separately and matched afterwards, the cross-participant comparison does not require every component to be present in every participant, and individual modes such as time can be relaxed during clustering. This makes the framework suited to the settings OP-MEG is opening up: heterogeneous clinical cohorts, variable sensor layouts and naturalistic paradigms in which responses cannot be time-locked across participants. Validation on larger cohorts, with simulated ground truth and trial-resolved tensors, will determine how far this extends beyond the well-characterised visuomotor responses examined here.

## Acknowledgements

We are grateful to the participants who took part in this study and to the clinical and technical teams who supported data collection. We acknowledge support from the NIHR University College London Hospitals Biomedical Research Centre, the Well-come Trust (307306/Z/23/Z), UKRI Frontier Research (EP/X023060/1), and the Engineering and Physical Sciences Research Council (EP/Z533191/1). DB is supported by a UKRI Frontier Research Grant (EP/X023060/1).

## Author Contributions

HW and AR contributed equally to study design, data analysis and interpretation, and preparation of the original manuscript. SM and SP contributed to data acquisition and carried out the OP-MEG recordings. MW, DB and TT contributed to interpretation of the findings and critical review and revision of the manuscript. UV and MS conceived and supervised the study, contributed to the development of the analytical approach and interpretation of the findings, and critically reviewed and revised the manuscript. All authors reviewed and approved the final manuscript.

## Data and Code Availability

The analysis pipeline (tensor construction, NN-CPD, permutation testing and constrained consensus clustering) is available at https://github.com/hug0-w/OPM-MEG-cp-pipeline/tree/main. The OP-MEG recordings cannot be shared publicly because they are patient data; requests for access should be directed to the corresponding author.

## Appendix

**Table A1:** Participant demographics, trial counts and recording lengths. All participants were right handed.

| Participant | Age Range | Sex | Valid Trials | Recording length (min) |
|---|---|---|---|---|
| 1 | 35-40 | Male | 193 | 22 |
| 2 | 45-50 | Male | 198 | 10 |
| 3 | 40-50 | Female | 100 | 8 |

**Table A2:** Cortical regions of interest selected from the AAL2 atlas (70 regions, 35 bilateral pairs).

| No. | Left hemisphere | Right hemisphere |
|---|---|---|
| 1 | Precentral | Precentral |
| 2 | Frontal Sup 2 | Frontal Sup 2 |
| 3 | Frontal Mid 2 | Frontal Mid 2 |
| 4 | Frontal Inf Oper | Frontal Inf Oper |
| 5 | Frontal Inf Tri | Frontal Inf Tri |
| 6 | Frontal Inf Orb 2 | Frontal Inf Orb 2 |
| 7 | Rolandic Oper | Rolandic Oper |
| 8 | Supp Motor Area | Supp Motor Area |
| 9 | Frontal Sup Medial | Frontal Sup Medial |
| 10 | Frontal Med Orb | Frontal Med Orb |
| 11 | Rectus | Rectus |
| 12 | OFCmed | OFCmed |
| 13 | OFCant | OFCant |
| 14 | OFCpost | OFCpost |
| 15 | OFClat | OFClat |
| 16 | Calcarine | Calcarine |
| 17 | Cuneus | Cuneus |
| 18 | Lingual | Lingual |
| 19 | Occipital Sup | Occipital Sup |
| 20 | Occipital Mid | Occipital Mid |
| 21 | Occipital Inf | Occipital Inf |
| 22 | Fusiform | Fusiform |
| 23 | Postcentral | Postcentral |
| 24 | Parietal Sup | Parietal Sup |
| 25 | Parietal Inf | Parietal Inf |
| 26 | SupraMarginal | SupraMarginal |
| 27 | Angular | Angular |
| 28 | Precuneus | Precuneus |
| 29 | Paracentral Lobule | Paracentral Lobule |
| 30 | Heschl | Heschl |
| 31 | Temporal Sup | Temporal Sup |
| 32 | Temporal Pole Sup | Temporal Pole Sup |
| 33 | Temporal Mid | Temporal Mid |
| 34 | Temporal Pole Mid | Temporal Pole Mid |
| 35 | Temporal Inf | Temporal Inf |

**Figure A1:**
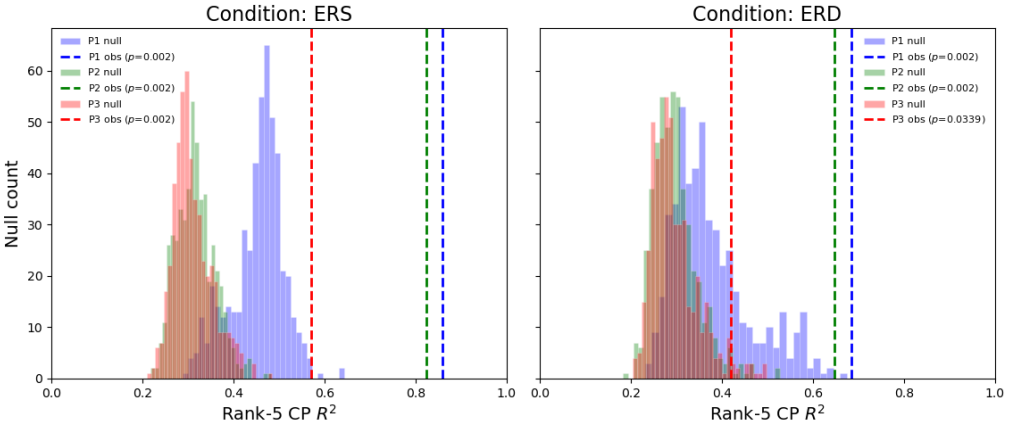
Permutation-based assessment of task-locked low-rank structure. Observed variance explained is compared with the null distribution obtained after independently circularly shifting trial time courses before tensor construction and decomposition.

### Rank Selection

Selecting the rank *R* is NP-hard (Hillar and Lim, 2013) and tensor-rank problems are computationally difficult. Model rank selection relies on empirical heuristics such as reconstruction error, CORCONDIA (Bro and Kiers, 2003) and stability (Williams et al., 2018). CORCONDIA assesses how far the fitted CP solution departs from an ideal super diagonal core: non-CP structure distributes mass into the off-diagonal core elements, lowering the diagnostic (Bro and Kiers, 2003). It was not used here, as the metric degrades for the highly collinear factors typical of non-negative decompositions, yielding unreliable estimates. We therefore used stability and variance explained computed using reconstruction error. For stability, the ALS NN-CPD was run *D* = 30 times from random initialisations at each rank; all runs were compared against the lowest-error reference using the model-similarity score of (Williams et al., 2018) originally adapted from (Tomasi and Bro, 2006), with the Hungarian algorithm used to resolve CPD’s permutation invariance (Appendix: Algorithm 1). Scores were averaged across runs into a single stability value per rank (1.0 = identical runs). Stability falls at higher ranks as the model fits noise and reproducibility drops. Variance explained provides an elbow criterion: the optimal rank is where additional components cease to explain meaningful variance and begin splitting existing ones. Although high ranks can retain broad similarity even as stability declines (Williams et al., 2018), a conservative choice favouring reproducibility was made (e.g. *R* = 3; Figure A2).

**Figure A2:**
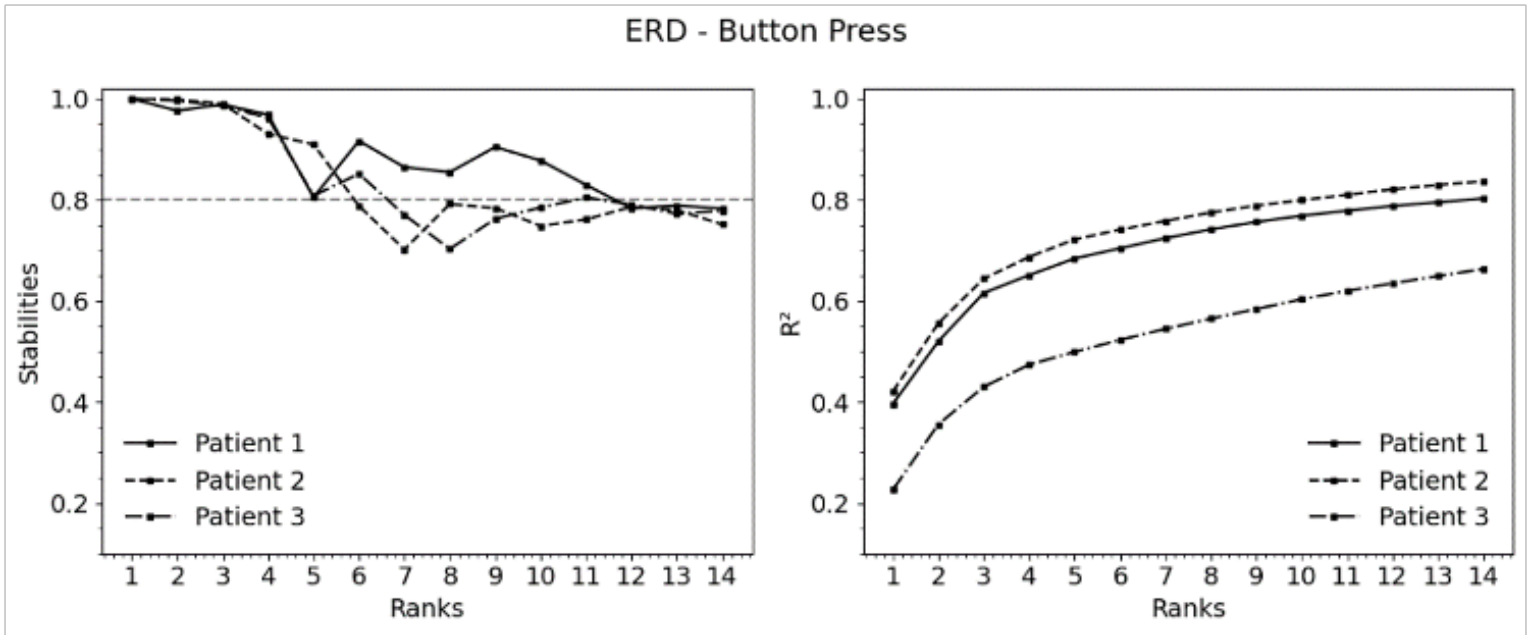
Rank-selection diagnostics for the ERD tensors. Reconstruction performance and stability across 30 random initialisations are shown as a function of rank. The selected rank balanced additional variance explained against reproducibility across initialisations. The ERS tensors showed the same trend but with a slightly larger decrease in stability for *R >* 3 across all participants.

For all participants ERS and ERD tensors, *R* = 3 was chosen as the highest stable rank across all participants roughly at the elbow of the variance plots.

#### Algorithm 1

Similarity Score

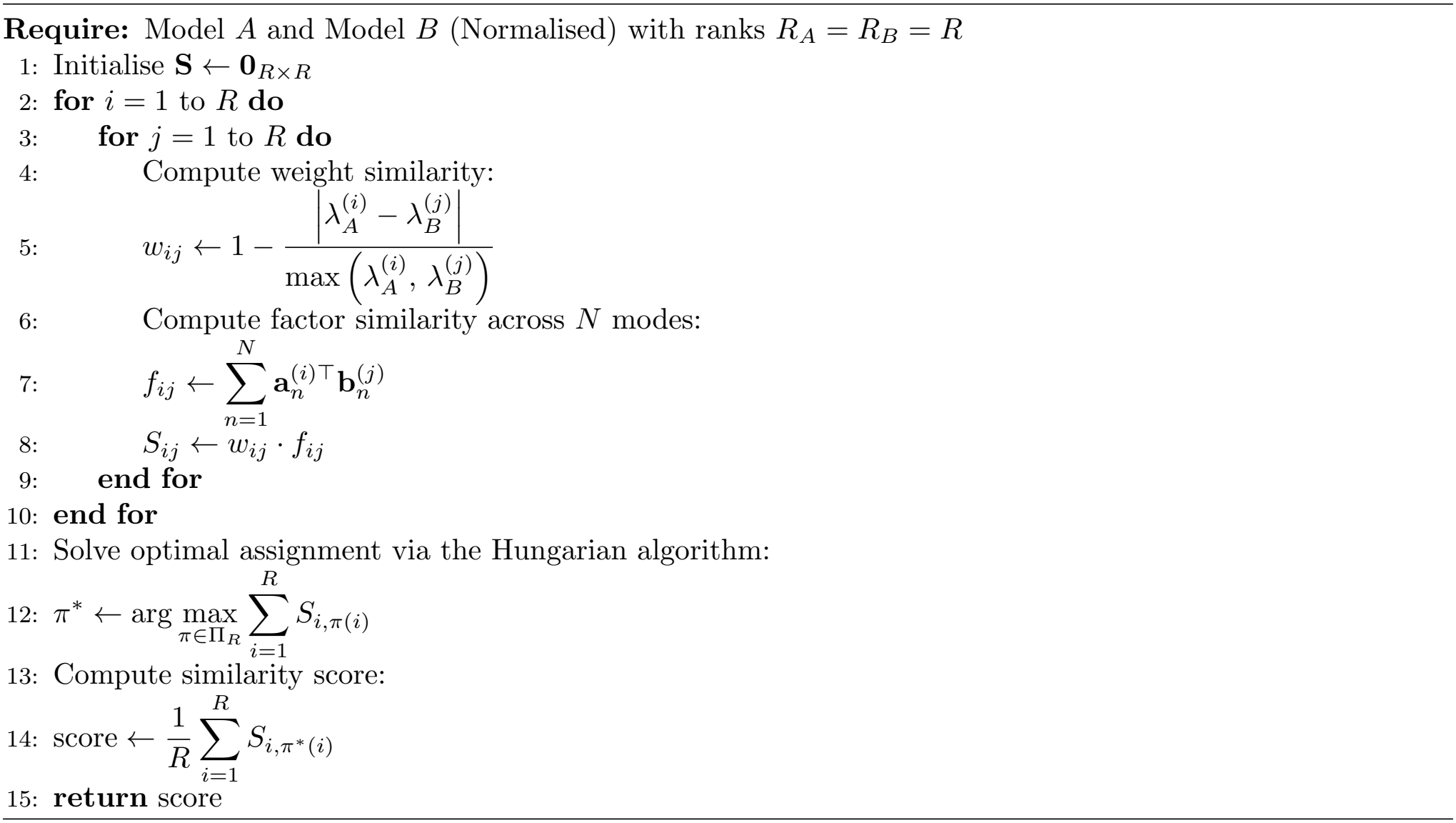

#### Other Findings

A third cluster was common to participants 2 and 3, dominated by gamma (*>* 30 Hz) synchronisation. Its temporal signature fluctuated quasi-periodically at ~5–10 Hz from ~200 ms after the button press. Spatially, the cluster was less consistent: the participant 2 component loaded mainly on visual cortex, whereas the participant 3 component loaded on frontal regions, and its spectral signature additionally carried theta power (5–8 Hz). The periodic gamma envelope is consistent with phase–amplitude coupling (PAC) between low-frequency phase and gamma amplitude. Alpha–gamma PAC has been reported over posterior cortex at rest (Osipova et al., 2008), increases there during visual tasks (Voytek et al., 2010), and has been interpreted as pulsed inhibition implementing top-down gating (Bonnefond and Jensen, 2015). Anteriorly, gamma couples preferentially to theta rather than alpha (Voytek et al., 2010), and frontotemporal theta–gamma coupling has been linked to working-memory maintenance (Daume et al., 2024). The frontal loading in participant 3 resembles this pattern, although co-modulated theta power does not itself indicate phase coupling. Because the tensor contains trial-averaged power, the observed periodicity requires the low-frequency phase to be reset by the event; confirming PAC would require a direct coupling measure on the source time series.

**Figure A3:**
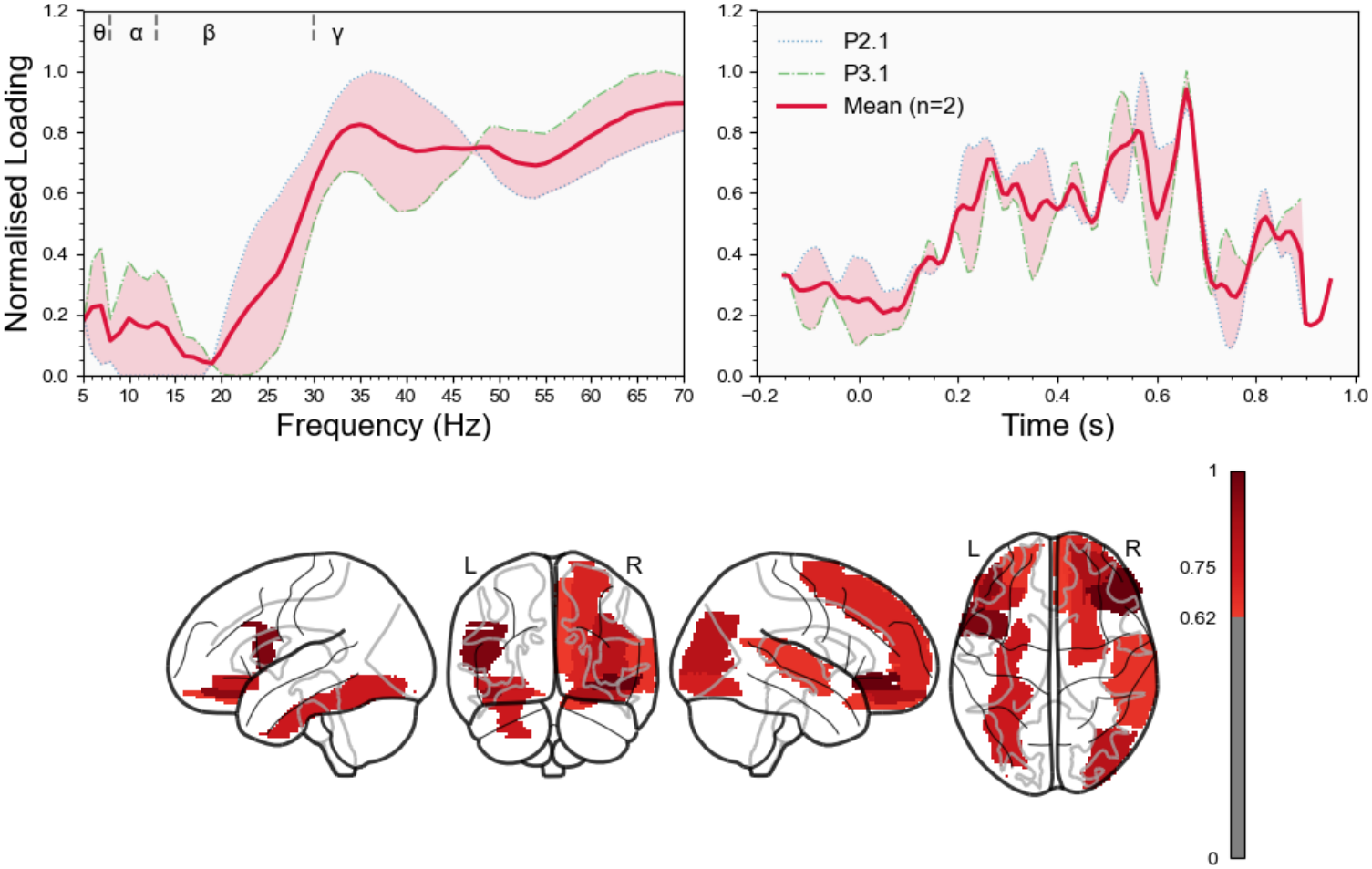
Additional ERS cluster identified in participants 2 and 3, showing gamma-band loadings and rhythmic fluctuations in the temporal loading profiles.

